# Duplicates, redundancies, and inconsistencies in the primary nucleotide databases: a descriptive study

**DOI:** 10.1101/085019

**Authors:** Qingyu Chen, Justin Zobel, Karin Verspoor

## Abstract

GenBank, the EMBL European Nucleotide Archive, and the DNA DataBank of Japan, known collectively as the International Nucleotide Sequence Database Collaboration or INSDC, are the three most significant nucleotide sequence databases. Their records are derived from laboratory work undertaken by different individuals, by different teams, with a range of technologies and assumptions, and over a period of decades. As a consequence, they contain a great many duplicates, redundancies, and inconsistencies, but neither the prevalence nor the characteristics of various types of duplicates have been rigorously assessed. Existing duplicate detection methods in bioinformatics only address specific duplicate types, with inconsistent assumptions; and the impact of duplicates in bioinformatics databases has not been carefully assessed, making it difficult to judge the value of such methods. Our goal is to assess the scale, kinds, and impact of duplicates in bioinformatics databases, through a retrospective analysis of merged groups in INSDC databases. Our outcomes are threefold: (1) We analyse a benchmark dataset consisting of duplicates manually identified in INSDC – a dataset of 67,888 merged groups with 111,823 duplicate pairs across 21 organisms from INSDC databases – in terms of the prevalence, types, and impacts of duplicates. (2) We categorise duplicates at both sequence and annotation level, with supporting quantitative statistics, showing that different organisms have different prevalence of distinct kinds of duplicate. (3) We show that the presence of duplicates has practical impact via a simple case study on duplicates, in terms of GC content and melting temperature. We demonstrate that duplicates not only introduce redundancy, but can lead to inconsistent results for certain tasks. Our findings lead to a better understanding of the problem of duplication in biological databases.

## 1. Introduction

Many kinds of database contain multiple instances of records. These instances may be identical, or may be similar but with inconsistencies; in traditional database contexts, this means that the same entity may be described in conflicting ways. In this paper, as elsewhere in the literature, we refer to such repetitions – whether redundant or inconsistent – as *duplicates*. The presence of any of these kinds of duplicate has the potential to confound analysis that aggregates or reasons from the data. Thus it is valuable to understand the extent and kind of duplication, and to have methods for managing it.

We regard two records as duplicates if, in the context of a particular task, the presence of one means that the other is not required. Duplicates are an ongoing data quality problem reported in diverse domains, including business (1), health care (2), and molecular biology (3). The five most severe data quality issues in general domains have been identified as redundancy, inconsistency, inaccuracy, incompleteness, and untimeliness (4). We must consider whether these issues also occur in nucleotide sequence databases.

GenBank, the EMBL European Nucleotide Archive (ENA), and the DNA DataBank of Japan (DDBJ), the three most significant nucleotide sequence databases, together form the International Nucleotide Sequence Database Collaboration (INSDC) (5). The problem of duplication in the bioinformatics domain is in some respects more acute than in general databases, as the underlying entities being modelled are imperfectly defined, and scientific understanding of them is changing over time. As early as 1996, data quality problems in sequence databases were observed, and concerns were raised that these errors may affect the interpretation (6). However, data quality problems persist, and current strategies for cleansing do not scale (7). Technological advances have led to rapid generation of genomic data. Data is exchanged between repositories that have different standards for inclusion.Ontologies are changing over time, as are data generation and validation methodologies. Data from different individual organisms, with genomic variations, may be conflated, while some data that is apparently duplicated – such as identical sequences from different individuals, or even different species – may in fact not be redundant at all. The same gene may be stored multiple times with flanking regions of different length, or, more perniciously, with different annotations. In the absence of a thorough study of the prevalence and kind of such issues, it is not known what impact they might have in practical biological investigations.

A range of duplicate detection methods for biological databases have been proposed (8-18). However, this existing work has defined duplicates in inconsistent ways, usually in the context of a specific method for duplicate detection. For example, some define duplicates solely on the basis of gene sequence identity, while others also consider metadata. These studies addressed only some of the kinds of duplication, and neither the prevalence nor the characteristics of different kinds of duplicate were measured.

A further, fundamental issue is that duplication (redundancy or inconsistency) cannot be defined purely in terms of the content of a database. A pair of records might only be regarded as duplicates in the context of a particular application. For example, two records that report the coding sequence for a protein may be redundant for tasks that concern RNA expression, but not redundant for tasks that seek to identify their (different) locations in the genome. Methods that seek to de-duplicate databases based on specific assumptions about how the data is to be used will have unquantified, potentially deleterious, impact on other uses of the same data.

Thus definitions of duplicates, redundancy, and inconsistency depend on context. In standard databases, a duplicate occurs when a unique entity is represented multiple times. In bioinformatics databases, duplicates have different representations, and the definition of ‘entity’ may be unclear. Also, duplicates arise in a variety of ways. The same data can be submitted by different research groups to a database multiple times, or to different databases without cross-reference. An updated version of a record can be entered while the old version still remains. Or there may be records representing the same entity, but with different sequences or different annotations.

Duplication can affect use of INSDC databases in a variety of ways. A simple example is that redundancy (such as records with near-identical sequences and consistent annotations) creates inefficiency, both in automatic processes such as search, and in manual assessment of the results of search.

More significantly, sequences or annotations that are inconsistent can affect analyses such as quantification of the correlation between coding and non-coding sequences (19), or finding of repeat sequence markers (20). Inconsistencies in functional annotations (21) have the potential to be confusing; despite this, an assessment of 37 North American *branchiobdellidans* records concluded that nearly half are inconsistent with the latest taxonomy (22). Function assignments may rely on the assumption that similar sequences have similar function (23), but repeated sequences may bias the output sequences from the database searches (24).

### Why care about duplicates?

Research in other disciplines has emphasised the importance of studying duplicates. Here we assemble comments on the impacts of duplicates in biological databases, derived from public or published material and curator interviews:

1. ***Duplicates lead to redundancies:*** ‘Automated analyses contain a significant amount of redundant data and therefore violate the principles of normalization… In a typical Illumina Genomestudio results file 63% of the output file is composed of unnecessarily redundant data’ (25). ‘High redundancy led to an increase in the size of UniProtKB (TrEMBL), and thus to the amount of data to be processed internally and by our users, but also to repetitive results in BLAST searches … 46.9 million (redundant) entries were removed (in 2015).’^1^ We explain the TrEMBL redundancy issue in detail below.
2. ***Duplicates lead to inconsistencies:*** ‘Duplicated samples might provide a false sense of confidence in a result, which is in fact only supported by one experimental data point’ (26), ‘two genes are present in the duplicated syntenic regions, but not listed as duplicates (true duplicates but are not labelled). This might be due to local sequence rearrangements that can influence the results of global synteny analysis’ (25).
3. ***Duplicates waste curation effort and impair data quality:*** ‘for UniProtKB/SwissProt, as everything is checked manually, duplication has impacts in terms of curation time. For UniProtKB/TrEMBL, as it (duplication) is not manually curated, it will impact quality of the dataset’.^2^
4. ***Duplicates have propagated impacts*** even after being detected or removed: ‘Highlighting and resolving missing, duplicate or inconsistent fields … ∼20% of (these) errors require additional rebuild time and effort from both developer and biologist’ (27), ‘The removal of bacterial redundancy in UniProtKB (and normal flux in protein) would have meant that nearly all (>90%) of Pfam (a highly curated protein family database using UniProtKB data) seed alignments would have needed manual verification (and potential modification) …This imposes a significant manual biocuration burden’ (28).

The presence of duplicates is not always problematic, however. For instance, the purpose of the INSDC databases is mainly to archive nucleotide records. Arguably, duplicates are not a significant concern from an archival perspective; indeed the presence of a duplicate may indicate that a result has been reproduced and should be viewed as confident. That is, duplicates can be evidence for correctness. Recognition of such duplicates supports record linkage and helps researchers to verify their sequencing and annotation processes. However, there is an implicit assumption that those duplicates have been labelled accurately. Without labelling, those duplicates may confuse users, whether or not the records represent the same entities.

To summarise, the question of duplication is context-dependent, and its significance varies in these contexts: different biological databases, different biocuration processes, and different biological tasks. But it is clear that we should still be concerned about duplicates in INSDC. Over 95% of UniProtKB data are from INSDC and parts of UniProtKB are heavily curated; hence duplicates in INSDC would delay the curation time and waste curation effort in this case. Furthermore, its archival nature does not limit the potential uses of the data; other uses may be impacted by duplicates. Thus it remains important to understand the nature of duplication in INSDC.

In this paper, we analyse the scale, kind and impacts of duplicates in nucleotide databases, to seek better understanding of the problem of duplication. We focus on INSDC records that have been reported as duplicates by manual processes and then merged. As advised to us by database staff, submitters spot duplicates and are the major means of quality checking in these databases; duplicates are also reported by other users, and in some cases directly identified by curators. Revision histories of records track the merges of duplicates. Based on an investigation of the revision history, we collected and analysed 67,888 merged groups containing 111,823 duplicate pairs, across 21 major organisms. This is one of three benchmarks of duplicates that we have constructed (53). While it is the smallest and most narrowly defined of the three benchmarks, it allows us to investigate the nature of duplication in INSDC as it arises during generation and submission of biological sequences, and facilitates understanding the value of later curation.

Our analysis demonstrates that various duplicate types are present, and that their prevalence varies between organisms. We also consider how different duplicate types may impact biological studies. We provide a case study, an assessment of sequence GC content and of melting point, to demonstrate the potential impact of various kinds of duplicates. We show that the presence of duplicates can alter the results, and thus demonstrate the need for accurate recognition and management of duplicates in genomic databases.

## 2. Background

While the task of detecting duplicate records in biological databases has been explored, previous studies have made a range of inconsistent assumptions about duplicates. Here, we review and compare these prior studies.

### 2.1 Definitions of duplication

In the introduction, we described repeated, redundant, and inconsistent records as *duplicates*. We use a broad definition of duplicates because no precise technical definition will be valid in all contexts. ‘Duplicate’ is often used to mean that two (or more) records refer to the same entity, but this leads to two further definitional problems: determining what ‘entities’ are and what ‘same’ means. Considering a simple example, if two records have the same nucleotide sequences, are they duplicates? Some people may argue that they are, because they have exactly the same sequences, but others may disagree because they could come from different organisms.

These kinds of variation in perspective have led to a great deal of inconsistency. Table 1 shows a list of biological databases from 2009 to 2015 and their corresponding definitions of duplicates. We extracted the definition of duplicates, if clearly provided; alternatively, we interpreted the definition based on the examples of duplicates or other related descriptions from the database documentation. It can be observed that the definition dramatically varies between databases, even those in the same domain. Therefore we reflectively use a broader definition of duplicates rather than an explicit or narrow one. In this work we consider records that have been merged during a manual or semi-automatic review as duplicates. We explain the characteristics of the merged record dataset in detail later.

**Table 1.** Definitions of ‘duplicate’ in genomic databases from 2009 to 2015.

| Database | Domain | Interpretation of the term ‘duplicate’ |
| --- | --- | --- |
| (29) | biomolecular interaction network | repeated interactions between protein to protein, protein to DNA, gene to gene; same interactions but in different organism-specific files |
| (30) | gene annotation | (near) identical genes; fragments; incomplete gene duplication; and different stages of gene duplication |
| (31) | gene annotation | near or identical coding genes |
| (32) | gene annotation | same measurements on different tissues for gene expression |
| (33) | genome characterization | records with same meta data; same records with inconsistent meta data; same or inconsistent record submissions |
| (34) | genome characterization | create a new record with the configuration of a selected record |
| (35) | ligand for drug discovery | records with multiple synonyms; for example, same entries for TR4 (Testicular Receptor 4) but some used a synonym TAK1 (a shared name) rather than TR4 |
| (36) | peptidase cleavages | cleavages being mapped into wrong residues or sequences |

Databases in the same domain, for example gene annotation, may be specialized for different perspectives, such as annotations on genes in different organisms or different functions, but they arguably belong to the same broad domain.

A pragmatic definition for *duplication* is that a pair of records A and B are duplicates if the presence of A means that B is not required, that is, B is redundant in the context of a specific task or is superseded by A. This is, after all, the basis of much record merging, and encompasses many of the forms of duplicate we have observed in the literature. Such a definition provides a basis for exploring alternative technical definitions of what constitutes a duplicate and provides a conceptual basis for exploring duplicate detection mechanisms. We recognise that (counterintuitively) this definition is asymmetric, but it reflects the in-practice treatment of duplicates in the INSDC databases. We also recognize that the definition is imperfect, but the aim of our work is to establish a shared understanding of the problem, and it is our view that a definition of this kind provides a valuable first step.

### 2.2 Duplicates based on a simple similarity threshold (redundancies)

In some previous work, a single sequence similarity threshold is used to find duplicates (8,9,11,14,16,18). In this work, duplicates are typically defined as records with sequence similarity over a certain threshold, and other factors are not considered. These kinds of duplicates are often referred to as approximate duplicates or near duplicates (37), and are interchangeable with redundancies. For instance, one study located all records with over 90% mutual sequence identity (11). (A definition that allows efficient implementation, but is clearly poor from the point of view of the meaning of the data; an argument that 90% similar sequences are duplicated, but that 89% similar sequences are not, does not reflect biological reality.) A sequence identity threshold also applies in the CD-HIT method for sequence clustering, where it is assumed that duplicates have over 90% sequence identity (38). The sequence-based approach also forms the basis of the non-redundant database used for BLAST (39).

Methods based on the assumption that duplication is equivalent to high sequence similarity usually share two characteristics. First, efficiency is the highest priority; the goal is to handle large datasets. While some of these methods also consider sensitivity (40), efficiency is still the major concern. Second, in order to achieve efficiency, many methods apply heuristics to eliminate unnecessary pairwise comparisons. For example, CD-HIT estimates the sequence identity by word (short substring) counting and only applies sequence alignment if the pair is expected to have high identity.

However, duplication is not simply redundancy. Records with similar sequences are not necessarily duplicates and vice versa. As we will show later, some of the duplicates we study are records with close to exactly identical sequences, but other types also exist. Thus use of a simple similarity threshold may mistakenly merge distinct records with similar sequences (false positives) and likewise may fail to merge duplicates with different sequences (false negatives). Both are problematic in specific studies (41,42).

### 2.3 Duplicates based on expert labelling

A simple threshold can find only one kind of duplicate, while others are ignored. Previous work on duplicate detection has acknowledged that expert curation is the best strategy for determining duplicates, due to the rich experience, human intuition and the possibility of checking external resources that experts bring (43-45). Methods using human-generated labels aim to detect duplicates precisely, either to build models to mimic expert curation behaviour (44), or to use expert curated datasets to quantify method performance (46).They can find more diverse types than using a simple threshold, but are still not able to capture the diversity of duplication in biological databases. The prevalence and characteristics of each duplicate type are still not clear. This lack of identified scope introduces restrictions that, as we will demonstrate, impair duplicate detection.

Korning et al. (13) identified two types of duplicates: the same gene submitted multiple times (near-identical sequences), and different genes belonging to the same family. In the latter case, the authors argue that, since such genes are highly related, one of them is sufficient to represent the others. However, this assumption that only one version is required is task-dependent; as noted in the introduction, for other tasks the existence of multiple versions is significant. To our knowledge, this is the first published work that identified different kinds of duplicates in bioinformatics databases, but the impact, prevalence, and characteristics of the types of duplicates they identify is not discussed.

Koh et al. (12) separated the fields of each gene record, such as species and sequences, and measured the similarities among these fields. They then applied association rule mining to pairs of duplicates using the values of these fields as features. In this way, they characterized duplicates in terms of specific attributes and their combination. The classes of duplicates considered were broader than Korning et al.’s, but are primarily records containing the same sequence, specifically: (1) the same sequence submitted to different databases; (2) the same sequence submitted to the same database multiple times; (3) the same sequence with different annotations; and (4) partial records. This means that the (near-)identity of the sequence dominates the mined rules. Indeed, the top ten rules generated from Koh et al.’s analysis share the feature that the sequences have exact (100%) sequence identity.

This classification is also used in other work (10,15,17), which therefore has the same limitation. This work again does not consider the prevalence and characteristics of the various duplicate types. While Koh has a more detailed classification in her thesis (47), the problem of characterization of duplicates remains.

In this previous work, the potential impact on bioinformatics analysis caused by duplicates in gene databases is not quantified. Many refer to the work of Muller et al. (7) on data quality, but Muller et al. do not encourage the study of duplicates; indeed, they claim that duplicates do not interfere with interpretation, and even suggest that duplicates may in fact have a positive impact, by ‘providing evidence of correctness’. However, the paper does not provide definitions or examples of duplicates, nor does it provide case studies to justify these claims.

### 2.4 Duplication persists due to its complexity

De-duplication is a key early step in standard data curation. Amongst biological databases, UniProt databases are well-known to have high quality data and detailed curation processes (48).We find that they have four different assumptions: ‘one record for 100% identical full-length sequences in one species’; ‘one record per gene in one species’; ‘one record for 100% identical sequences over the entire length, regardless of the species’; and ‘one record for 100% identical sequences, including fragments, regardless of the species’, for UniProtKB/TrEMBL, UniProtKB/SwissProt, UniParc, and UniRef100 respectively.^3^ We note the emphasis on sequence identity in these requirements.

Each database has its specific design and purpose, so the assumptions made about duplication differ. One community may consider a given pair to be a duplicate whereas other communities may not. The definition of duplication varies between biologists, database staff and computer scientists. In different curated biological databases, de-duplication is handled in different ways. It is far more complex than a simple similarity threshold; we want to analyse duplicates that are labelled based on human judgements rather than using a single threshold. Therefore, we created three benchmarks of nucleotide duplicates from different perspectives (53). In this work we focus on analysing one of these benchmarks, containing records directly merged in INSDC. Merging of records is a way to address data duplication. Examination of merged records facilitates understanding of what constitutes duplication.

Recently, in TrEMBL, UniProt staff observed that it had a high prevalence of redundancy. A typical example is that 1692 strains of Mycobacterium tuberculosis have been represented in 5.97 million entries, because strains of this same species have been sequenced and submitted multiple times. UniProt staff have expressed concern that such high redundancy will lead to repetitive results in BLAST searches. Hence they used a mix of manual and automatic approaches to de-duplicate bacterial proteome records, and removed 46.9 million entries in April 2015.^4^ A ‘duplicate’ proteome is selected by identifying: (a) two proteomes under the same taxonomic species group, (b) having over 90% identity, and (c) selecting the proteome of the pair with the highest number of similar proteomes for removal; specifically, all protein records in TrEMBL belonging to the proteome will be removed.^5^ If proteome A and B satisfy criteria (a) and (b), and proteome A has 5 other proteomes with over 90% identity, whereas proteome B only has one, A will be removed rather than B. This notion of a duplicate differs from those above, emphasising the context dependency of the definition of a ‘duplicate’. This de-duplication strategy is incomplete as it removes only one kind of duplicate, and is limited in application to full proteome sequences; the accuracy and sensitivity of the strategy is unknown. Nevertheless removing one duplicate type already significantly reduces the size of TrEMBL. This not only benefits database search, but also affects studies or other databases using TrEMBL records.

This de-duplication is considered to be one of the two significant changes in UniProtKB database in 2015 (the other change being the establishment of a comprehensive reference proteome set) (28). It clearly illustrates that duplication in biological databases is not a fully solved problem and that de-duplication is necessary.

Overall, we can see that foundational work on the problem of duplication in biological sequence databases has not previously been undertaken. There is no prior thorough analysis of the presence, kind, and impact of duplicates in these databases.

## 3. Data and methods

Exploration of duplication and its impacts requires data. We have collected and analysed duplicates from INSDC databases to create a benchmark set, as we now discuss.

### 3.1 Collection of duplicates

Some of the duplicates in INSDC databases have been found and then merged into one representative record. We call this record the *exemplar*, that is, the current record retained as a proxy for a set of records. Staff working at EMBL ENA advised us (by personal communication) that a merge may be initiated by original record submitter, database staff, or occasionally in other ways. We further explain the characteristics of the merged dataset below, but note that records are merged for different reasons, showing that diverse causes can lead to duplication. The merged records are documented in the revision history. For instance, GenBank record `gi:6017069` is the complete sequence of both BACR01G10 and BACR05I08 clones for chromosome 2 in *Drosophila melanogaster*. Its revision history^6^ shows that it has replaced two records `gi:6015178` and `gi:6012087`, because they are ‘SEQUENCING IN PROGRESS’ records with 57 and 21 unordered pieces for BACR01G10 and BACR05I08 clones respectively. As explained in the supplementary materials, the groups of records can readily be fetched using NCBI tools.

For our analysis, we collected 67,888 groups (during 15–27 July 2015), which contained 111,823 duplicates (a given group can contain more than one record merge) across the 21 popular organisms used in molecular research listed in the NCBI Taxonomy web page.^7^ The data collection is summarized in Supplementary Table S1, and, the details of the collection procedure underlying the data are elaborated in the Supplementary file *Details of the record collection procedure.* As an example, the *Xenopus laevis* organism has 35,544 directly related records. Of these, 1,690 have merged accession IDs; 1,620 merged groups for 1,660 duplicate pairs can be identified in the revision history.

### 3.2 Characteristics of the duplicate collection

As explained in Section 2, we use a broad definition of duplicates. This data collection reflects the broad definition, and in our view is representative of an aspect of duplication: these are records that are regarded as similar or related enough to merit removal, that is, are redundant. The records were merged for different reasons, including:

- Changes to data submission policies. Before 2003, the sequence submission length limit was 350 kb. After releasing the limit, the shorter sequence submissions were merged into a single comprehensive sequence record.
- Updates of sequencing projects. Research groups may deposit current draft records; later records will merge the earlier ones.
- Merges from other data sources. For example, RefSeq uses INSDC records as a main source for genome assembly (49). The assembly is made according to different organism models and updated periodically and the records may be merged or split during each update (50).
- Merges by record submitters or database staff occur when they notice multiple submissions of the same record.

While the records were merged due to different reasons, they can all be considered duplicates. The various reasons for merging records represent the diversity. If those records above had not been merged, they would cause data redundancy and inconsistency.

These merged records are illustrations of the problem of duplicates rather than current instances to be cleaned. Once the records are merged, they are no longer active or directly available to database users. However, the obsolete records are still of value. For example, even though over 45 million duplicate records were removed from UniProt, the key database staff who were involved in this activity^8^ are still interested in investigating their characteristics. They would like to understand the similarity of duplicates for more rapid and accurate duplicate identification in future, and to understand their impacts, such as how their removal affects database search.

From the perspective of a submitter, those records removed from UniProtKB may not be duplicates, since they may represent different entities, have different annotations, and serve different applications. However, from a database perspective, they challenge database storage, searches, and curation (48). ‘Most of the growth in sequences is due to the increased submission of complete genomes to the nucleotide sequence databases’ (48). This also indicates that records in one data source may not be considered as duplicates, but do impact other data sources.

To our knowledge, our collection is the largest set of duplicate records merged in INSDC considered to date. Note that we have collected even larger datasets based on other strategies, including expert and automatic curation (51). We focus on this collection here, to analyse how submitters understand duplicates as one perspective. This duplicate dataset is based on duplicates identified by those closest to the data itself, the original data submitters, and is therefore of high quality.

We acknowledge that the data set is by its nature incomplete; the number of duplicates that we have collected is likely to be a vast undercounting of the exact or real prevalence of duplicates in the INSDC databases. There are various reasons for this that we detail here.

First, as mentioned above, both database staff and submitters can request merges. However, for submitters, records can only be modified or updated if they are the record owner. Other parties who want to update records that they did not themselves submit must get permission from at least one original submitter.^9^ In EMBL ENA, it is suggested to contact the original submitter first, but there is an additional process for reporting errors to the database staff.^10^ Due to the effort required for these procedures, the probability that there are duplicates that have not been merged or labelled is very high.

Additionally, as the documentation shows, submitter-based updates or correction are the main quality control mechanisms in these databases. Hence the full collection of duplicates listed in Supplementary Table S1 presented in this work are limited to those identified by (some) submitters. Our other duplicate benchmarks, derived from mapping INSDC to Swiss-Prot and TrEMBL, contain many more duplicates (53). This implies that many more potential duplicates remain in INSDC.

The impact of curation on marking of duplicates can be observed in some organisms. The total number of records in *Bos taurus* is about 14% and 1.9% of the number of records in *Mus musculus* and *Homo sapiens*, respectively, yet *Bos taurus* has a disproportionately high number of duplicates in the benchmark: more than 20,000 duplicate pairs, which is close (in absolute terms) to the number of duplicates identified in the other two species. Another example is *Schizosaccharomyces pombe*, which only has around 4000 records but a relatively large number (545) of duplicate pairs have been found.

An organism may have many more duplicates if its sub-organisms are considered. The records counted in the table are directly associated to the listed organism; we did not include records belonging to any sub-organisms in this study. An example of the impact of this is record `gi:56384585`, which replaced 500 records in 2004.^11^ This record belongs to *Escherichia coli O157:H7 strain EDL933*, which is not directly associated to *Escherichia coli* and therefore not counted here. The collection statistics also demonstrate that 13 organisms contain at least some merged records for which the original records have different submitters. This is particularly evident in *Caenorhabditis elegans* and *Schizosaccharomyces pombe* (where 92.4% and 81.8%, respectively, of duplicate records are from different submitters). The possible explanations are that database staff merged those records, or that an error was reported to database staff by a third party.

This benchmark is the only resource currently available for duplicates directly merged in INSDC. Staff have also advised that there is currently no automatic process for collecting such duplicates.

### 3.3 Categorization of duplicates

Observing the duplicates in the collection, we find that some of them share the same sequences, whereas others have sequences with varied lengths. Some have been annotated by submitters with notes such as ‘WORKING DRAFT’. We therefore categorized records at both sequence level and annotation level. For sequence level, we identified five categories: *Exact sequences, Similar sequences, Exact fragments, Similar fragments,* and *Low-identity sequences*. For annotation level, we identified three categories: *Working draft, Sequencing-in-progress, and Predicted*. We do not restrict a duplicate instance to be in only one category.

This categorization represents diverse types of duplicates in nucleotide databases, and each distinct kind has different characteristics. As discussed previously, there is no existing categorization of duplicates with supporting measures or quantities in prior work. Hence we adopt this categorization and quantify the prevalence and characteristics of each kind, as a starting point for understanding the nature of duplicates in INSDC databases more deeply.

The detailed criteria and description of each category are as follows. For sequence level, we measured *local sequence identity* using BLAST (9). This measures whether two sequences share similar subsequences. We also calculated the *local alignment proportion* (the number of identical bases in BLAST divided by the length of the longer sequence of the pair) to estimate the possible coverage of the pair globally without performing a complete (expensive) global alignment. Details, including formulas, are provided in the supplementary materials *Details of measuring submitter similarity* and *Details of measuring sequence similarities*.

**Category 1, sequence level:** *Exact sequences*. This category consists of records that share exact sequences. We require that the local identity and local alignment proportion must both be 100%. While this cannot guarantee that the two sequences are exactly identical without a full global alignment, having both local identity and alignment coverage of 100% strongly implies that two records have the same sequences.

**Category 2, sequence level:** *Similar sequences*. This category consists of records that have near-identical sequences, where the local identity and local alignment proportion are less than 100% but no less than 90%.

**Category 3, sequence level:** *Exact fragments.* This category consists of records that have identical subsequences, where the local identity is 100% and the alignment proportion is less than 90%, implying that the duplicate is identical to a fragment of its replacement.

**Category 4, sequence level:** *Similar fragments*. By correspondence with the relationship between Categories 1 and 2, this category relaxes the constraints of Category 3. It has the same criteria of alignment proportion as Category 3, but reduces the requirement for local identity to no less than 90%.

**Category 5, sequence level:** *Low-identity sequences*. This category corresponds to duplicate pairs that exhibit weak or no sequence similarity. This category has three tests: first, the local sequence identity is less than 90%; second, BLAST output is ‘NO HIT’, that is, no significant similarity has been found; third, the expected value of the BLAST score is greater than 0.001, that is, the found match is not significant enough.

**Categories based on annotations** The categories at the annotation level are identified based on record submitters’ annotations in the ‘DEFINITION’ field. Some annotations are consistently used across the organisms, so we used them to categorise records.

If at least one record of the pair contains the words ‘WORKING DRAFT’, it will be classified as *Working draft*, and similarly for *Sequencing-in-progress* and *Predicted*, containing ‘SEQUENCING IN PROGRESS’ and ‘PREDICTED’, respectively.

A more detailed categorization could be developed based on this information. For instance, there are cases where both a duplicate and its replacement are working drafts, and other cases where the duplicate is a working draft while the replacement is the finalized record. It might also be appropriate to merge *Working draft* and *Sequencing-in-progress* into one category, since they seem to capture the same meaning. However, to respect the original distinctions made by submitters, we have retained it.

## 4. Presence of different duplicate types

Supplementary Table S2 shows the distribution of duplicates across our categories. Table 1 extracts a representative subset.

Recall that existing work mainly focuses on duplicates with similar or identical sequences. However, based on the duplicates in our collection, we observe that duplicates under the *Exact sequence* and *Similar sequence* categories only represent a fraction of the known duplicates. Only nine of the 21 organisms have *Exact sequence* as the most common duplicate type, and six organisms have small numbers of this type. Thus the general applicability of prior proposals for identifying duplicates is questionable.

Additionally, it is apparent that the prevalence of duplicate types is different across the organisms. For sequence-based categorization, for nine organisms the highest prevalence is Exact sequence (as mentioned above), for two organisms it is *Similar sequences*, for eight organisms it is *Exact fragments*, and for three organisms it is *Similar fragments* (one organism has been counted twice since *Exact sequence* and *Similar fragments* have the same count). It also shows that ten organisms have duplicates that have relatively low sequence identity.

Overall, even this simple initial categorization illustrates the diversity and complexity of known duplicates in the primary nucleotide databases. In other work (52), we reproduced a representative duplicate detection method using association rule mining (12) and evaluated it with a sample of 3498 merged groups from *Homo Sapiens*. The performance of this method was extremely poor. The major underlying issues were that the original dataset only contains duplicates with identical sequences and that the method did not consider diverse duplicate types.

Thus it is necessary to categorize and quantify duplicates to find out distinct characteristics held by different categories and organisms; we suggest that these different duplicate types must be separately addressed in any duplicate detection strategy.

## 5. Impacts of duplicates: case study

An interesting question is whether duplicates affect biological studies, and to what extent. As a preliminary investigation, we conducted a case study on two characteristics of DNA sequences: GC content and melting temperature. The GC content is the proportion of bases G and C over the sequence. Biologists have found that GC content is correlated with local rates of recombination in the human genome (53). The GC content of microorganisms is used to distinguish species during the taxonomic classification process.

The melting temperature of a DNA sequence is the temperature at which half of the molecules of the sequence form double strands, while another half are single-stranded, a key sequence property that is commonly used in molecular studies (54). Accurate prediction of the melting temperature is an important factor in experimental success (55). The GC content and the melting temperature are correlated, as the former is used in determination of the latter. The details of calculations of GC content and melting temperature are provided in the supplementary *Details of formulas in the case study*.

We computed and compared these two characteristics in two settings: by comparing exemplars with the original group, which contains the exemplars along with their duplicates; and by comparing exemplars with their corresponding duplicates, but with the exemplar removed.

Selected results are in Table 3 (visually represented in Figures 1 and 2) and 4 (visually represented in Figures 3 and 4) respectively (full results in Supplementary Tables S3 and S4). First, it is obvious that the existence of duplicates introduces much redundancy. After de-duplication, the size of original duplicate set is reduced by 50% or more for all the organisms shown in the table. This follows from the structure of the data collection.

**Figure 1.**
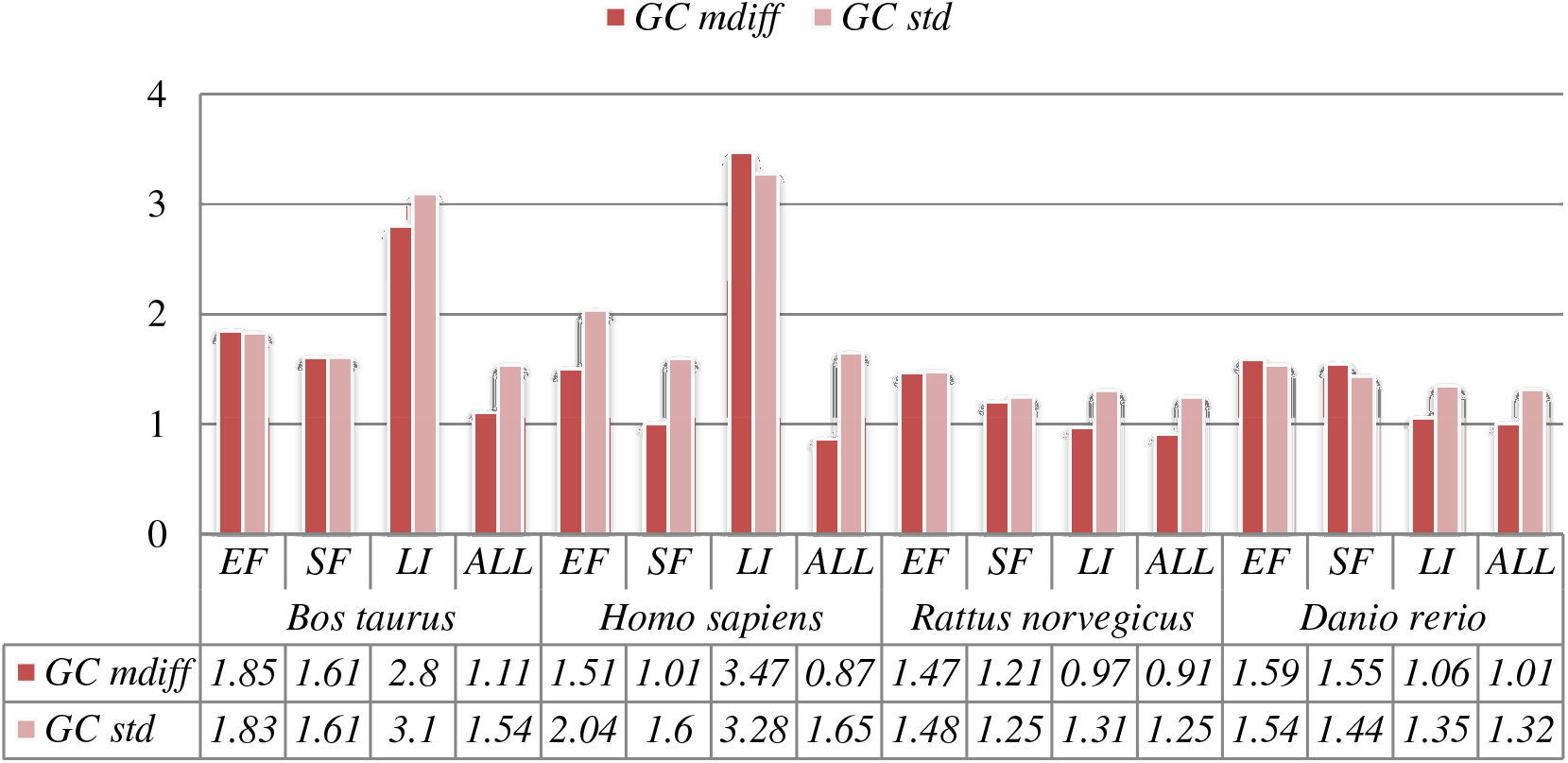
A selection of results for organisms in terms of GC content (Exemplar vs. Original merged groups) Categories are the same as Table 1; mdiff and std: the mean and standard deviation of absolute value of the difference between each exemplar and the mean of the original group respectively.

**Figure 2.**
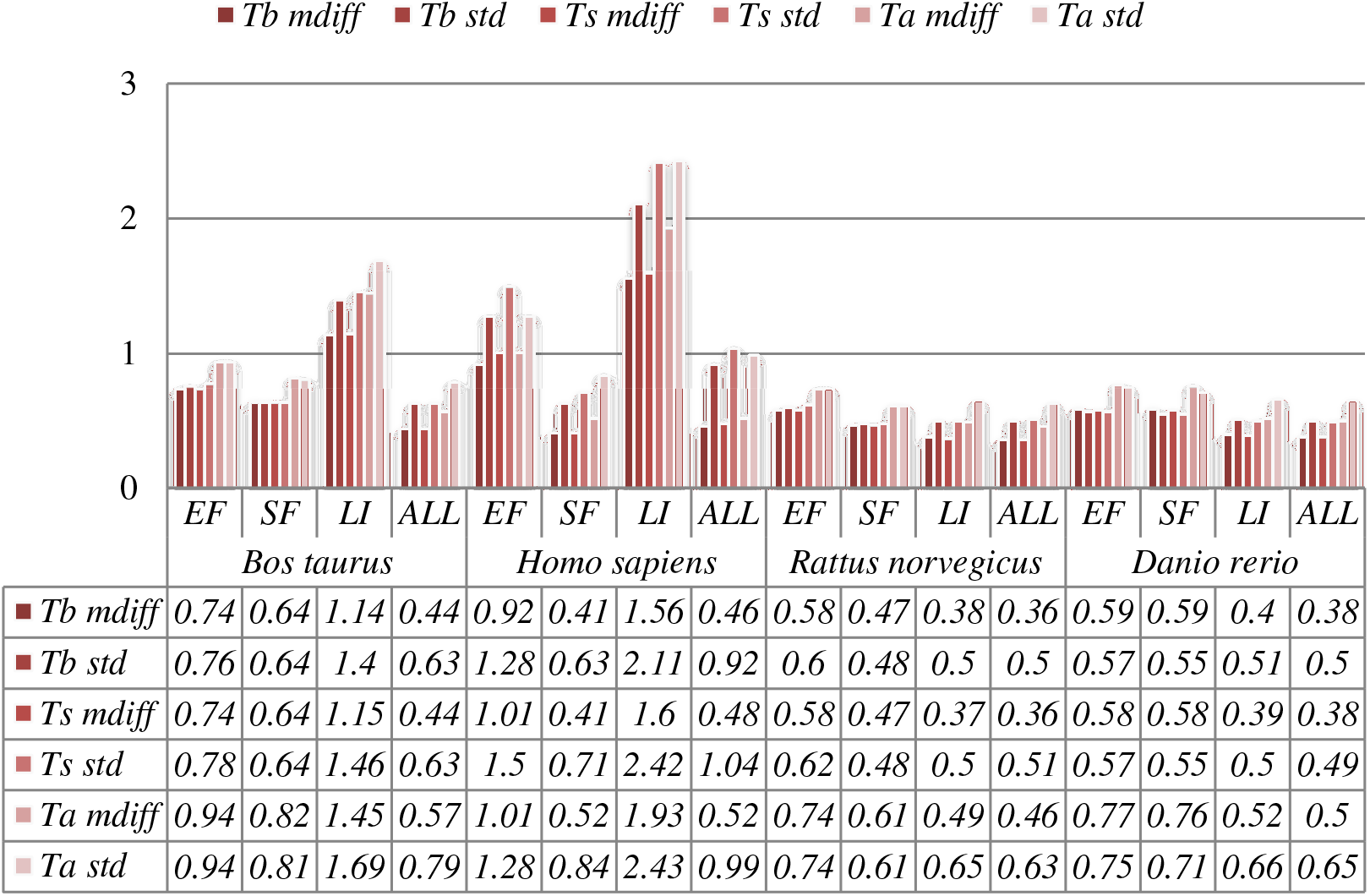
A selection of results for organisms in terms of melting temperatures (Exemplar vs. Original merged groups) mdiff and std: the mean and standard deviation of absolute value of the difference between each exemplar and the mean of the original group respectively; Tb, Ts, Ta: melting temperature calculated using basic, salted and advanced formula in supplement respectively.

**Figure 3.**
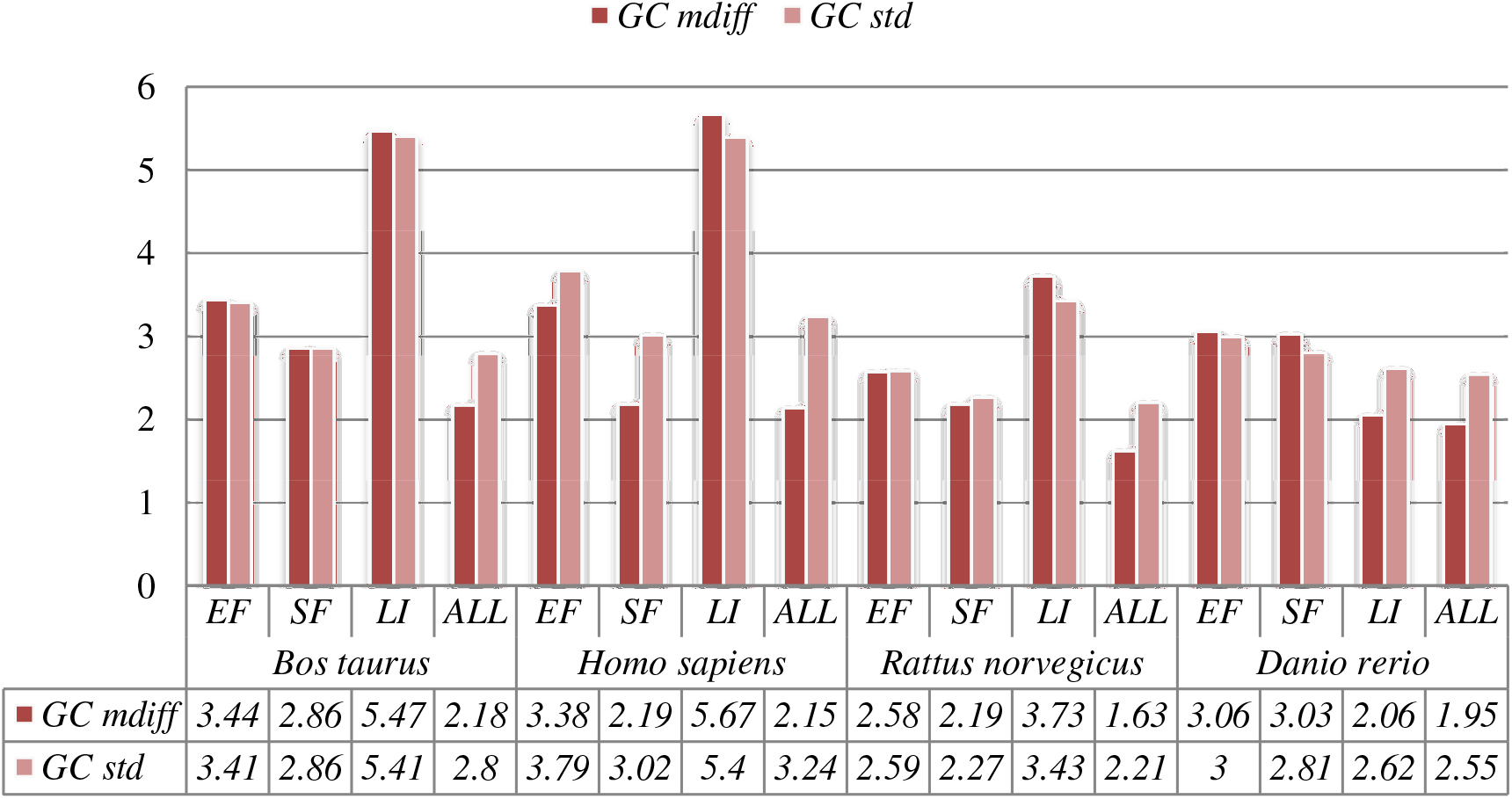
A selection of results for organisms in terms of GC content (Exemplar vs. Duplicate pairs) Categories are the same as Table 1; mdiff and std: the mean and standard deviation of absolute value of the difference between each exemplar and the mean of the duplicates group respectively.

**Figure 4.**
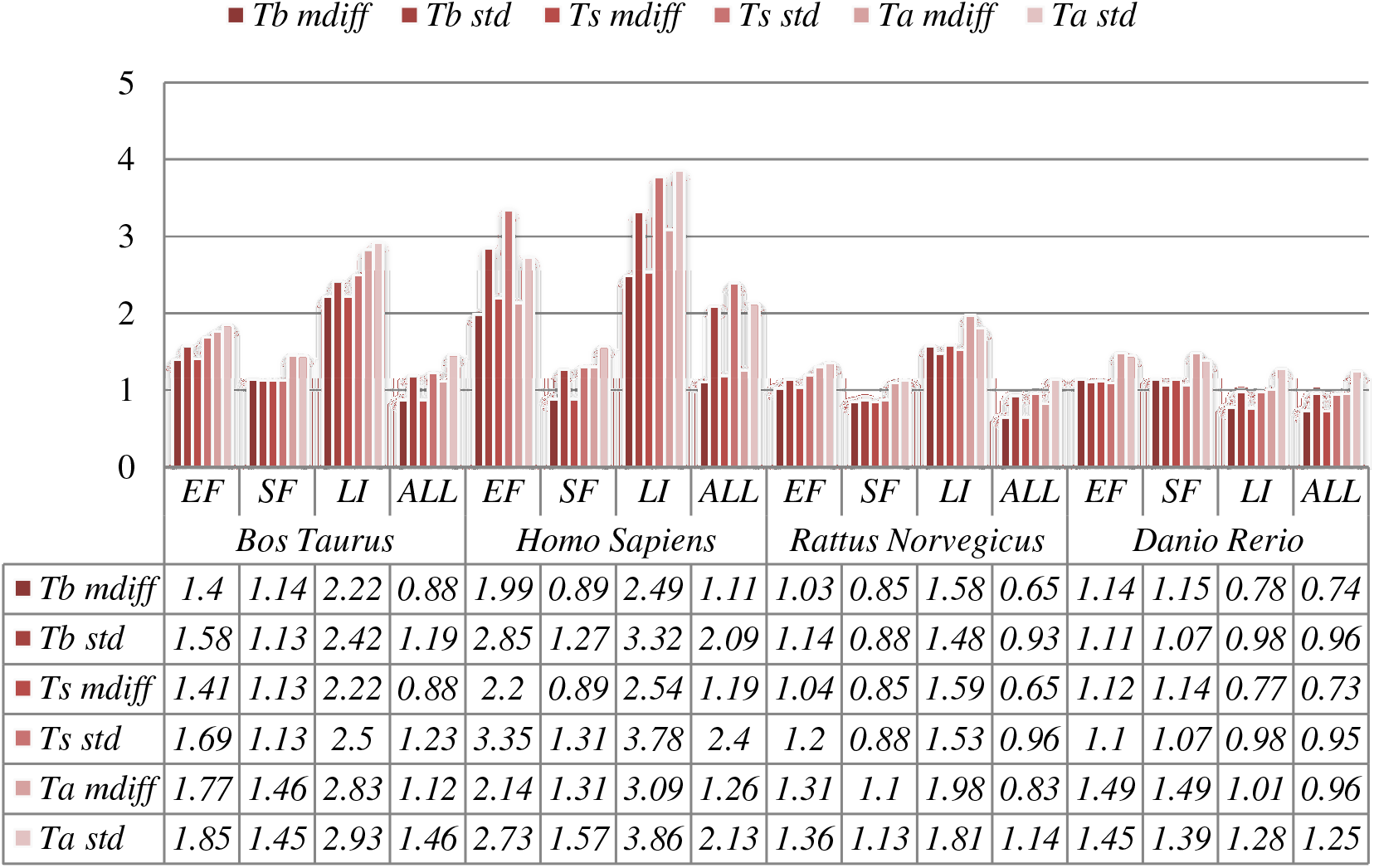
A selection of results for organisms in terms of melting temperatures (Exemplar vs. Duplicate pairs) mdiff and std: the mean and standard deviation of absolute value of the difference between each exemplar and the mean of the original group respectively; Tb, Ts, Ta: melting temperature calculated using basic, salted and advanced formula in supplement respectively.

Critically, it is also evident that all the categories of duplicates except *Exact sequences* introduce differences for the calculation of GC content and melting temperature. These *mdiff* (mean of difference) values are significant, as they exceed other experimental tolerances, as we explain below. (The values illustrating larger distinctions have been made bold in the table.) Table 2 already shows that exemplars have distinctions with their original groups. When examining exemplars with their specific pairs, the differences become even larger as shown in Table 3. Their mean differences and standard deviations are different, meaning that exemplars have distinct characteristics compared to their duplicates.

**Table 2.** Samples of duplicates types classified in both sequence level and annotation level

| Organism | Total records | Sequence-based |  |  |  |  | Annotation-based |  |  | Others |  |
| --- | --- | --- | --- | --- | --- | --- | --- | --- | --- | --- | --- |
|  |  | ES | SS | EF | SF | LI | WD | SP | PR | LS | UC |
| *Bos taurus* | 245,188 | 2,923 | 3,633 | 5,167 | 6,984 | 147 | 0 | 0 | 18,120 | 2,089 | 0 |
| *Homo sapiens* | 12,506,281 | 2,844 | 7,139 | 11,325 | 6,889 | 642 | 2,951 | 316 | 17,243 | 1,496 | 0 |
| *Caenorhabditis elegans* | 74,404 | 1736 | 7 | 109 | 44 | 5 | 0 | 121 | 0 | 0 | 0 |
| *Rattus norvegicus* | 318,577 | 2,511 | 5,302 | 7,556 | 3,817 | 107 | 0 | 0 | 15,382 | 2 | 0 |
| *Danio rerio* | 153,360 | 721 | 2740 | 1662 | 3504 | 75 | 1 | 34 | 7684 | 521 | 491 |
| *Mus musculus* | 1,730,941 | 2,597 | 4,689 | 6,678 | 7,377 | 379 | 1,926 | 1,305 | 16,510 | 2,011 | 1 |

Total records: Number of records in total directly belong to the organism (derived from NCBI taxonomy database);ES: exact sequences; SS: similar sequences; EF: exact fragments; SF: similar fragments; LI: low-identity sequences; WD: working draft; SP: sequencing-in-progress record; PR: predicted sequence; LS: long sequence; UC: unclassified pairs

**Table 3.**
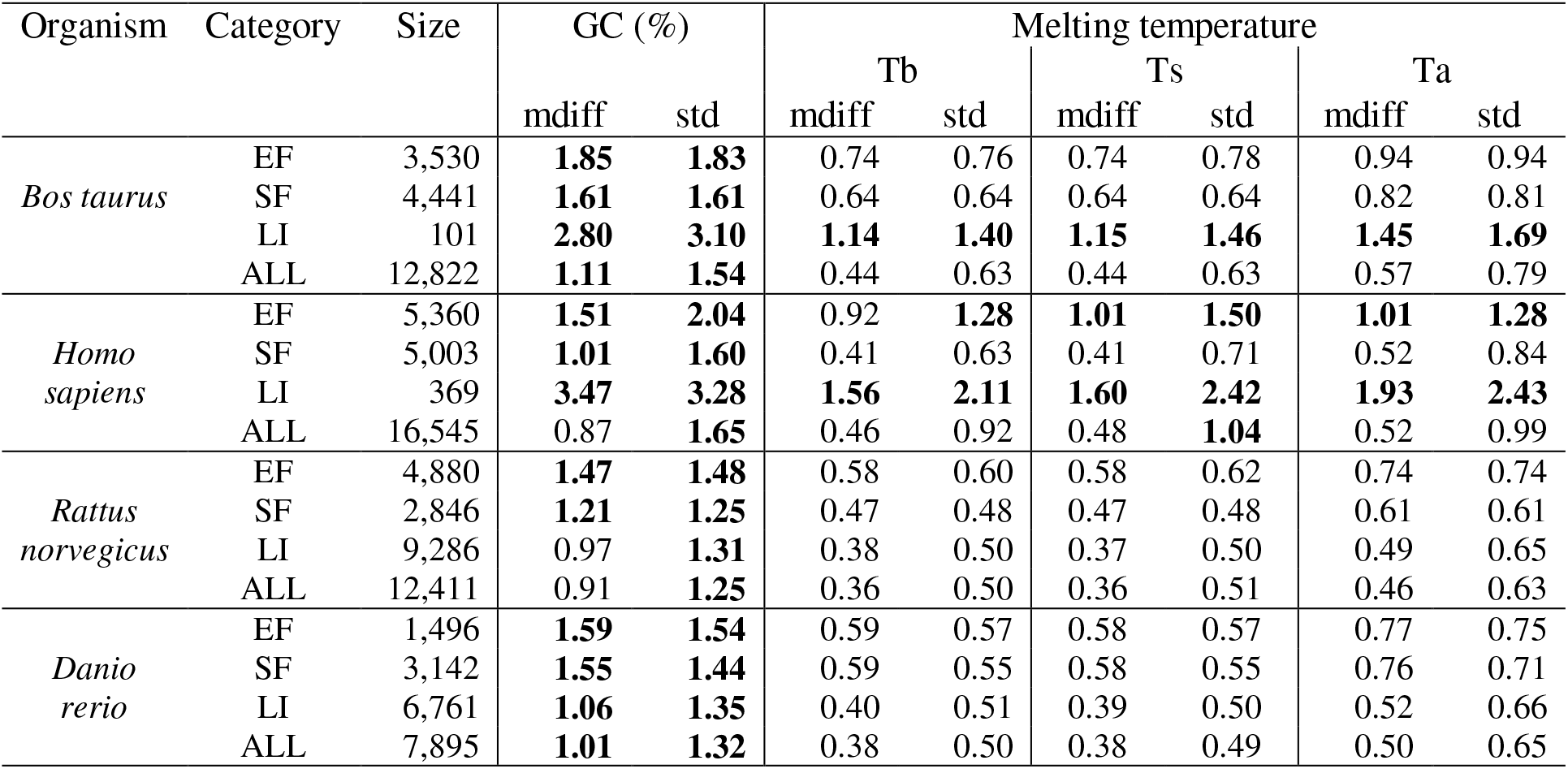
A selection of results for organisms in terms of GC content and melting temperatures (Exemplar vs. Original merged groups)

Categories are the same as Table 1; mdiff and std: the mean and standard deviation of absolute value of the difference between each exemplar and the mean of the original group respectively; Tb, Ts, Ta: melting temperature calculated using basic, salted and advanced formula in supplement respectively;

**Table 4.** A selection of results for organisms in terms of GC content and melting temperatures (Exemplar vs. Duplicate pairs)

| Organism | Category | Size | GC (%) |  | Melting temperature (°C) |  |  |  |  |  |
| --- | --- | --- | --- | --- | --- | --- | --- | --- | --- | --- |
|  |  |  | mdiff | std | Tb |  | Ts |  | Ta |  |
|  |  |  |  |  | mdiff | std | mdiff | std | mdiff | std |
| *Bos taurus* | EF | 5,167 | **3.44** | **3.41** | **1.40** | **1.58** | **1.41** | **1.69** | **1.77** | **1.85** |
|  | SF | 6,984 | **2.86** | **2.86** | **1.14** | **1.13** | **1.13** | **1.13** | **1.46** | **1.45** |
|  | LI | 149 | **5.47** | **5.41** | **2.22** | **2.42** | **2.22** | **2.50** | **2.83** | **2.93** |
|  | ALL | 20,945 | **2.18** | **2.80** | **0.88** | **1.19** | 0.88 | **1.23** | **1.12** | **1.46** |
| *Homo sapiens* | EF | 11,325 | **3.38** | **3.79** | **1.99** | **2.85** | **2.20** | **3.35** | **2.14** | **2.73** |
|  | SF | 6,890 | **2.19** | **3.02** | 0.89 | **1.27** | **0.89** | **1.31** | **1.31** | **1.57** |
|  | LI | 642 | **5.67** | **5.40** | **2.49** | **3.32** | **2.54** | **3.78** | **3.09** | **3.86** |
|  | ALL | 30,336 | **2.15** | **3.24** | **1.11** | **2.09** | **1.19** | **2.40** | **1.26** | **2.13** |
| *Rattus norvegicus* | EF | 7,556 | **2.58** | **2.59** | **1.03** | **1.14** | **1.04** | **1.20** | **1.31** | **1.36** |
|  | SF | 3,817 | **2.19** | **2.27** | 0.85 | 0.88 | 0.85 | 0.88 | **1.10** | **1.13** |
|  | LI | 107 | **3.73** | **3.43** | **1.58** | **1.48** | **1.59** | **1.53** | **1.98** | **1.81** |
|  | ALL | 19,295 | **1.63** | **2.21** | 0.65 | 0.93 | 0.65 | 0.96 | 0.83 | **1.14** |
| *Danio rerio* | EF | 1,662 | 3.06 | 3.00 | **1.14** | **1.11** | **1.12** | **1.10** | **1.49** | **1.45** |
|  | SF | 3,504 | 3.03 | 2.81 | **1.15** | **1.07** | **1.14** | **1.07** | **1.49** | **1.39** |
|  | LI | 7,684 | 2.06 | 2.62 | 0.78 | 0.98 | 0.77 | 0.98 | **1.01** | **1.28** |
|  | ALL | 9,227 | 1.95 | 2.55 | 0.74 | 0.96 | 0.73 | 0.95 | 0.96 | **1.25** |

Categories are the same as Table 1; mdiff and std: the mean and standard deviation of absolute value of the difference between each exemplar and the mean of the duplicates group respectively; Tb, Ts, Ta: melting temperature calculated using basic, salted and advanced formula in supplement respectively;

These differences are significant and can impact interpretation of the analysis. It has been argued in the context of a wet-lab experiment exploring GC content that well-defined species fall within a 3% range of variation in GC percentage (56). Here, duplicates under specific categories could introduce variation of close to or more than 3%. For melting temperatures, dimethyl sulfoxide (DMSO), an external chemical factor, is commonly used to facilitate the amplification process of determining the temperature. An additional 1% DMSO leads to a temperature difference ranging from 0.5°C to 0.75°C (54). However, six of our measurements in Homo sapiens have differences of over 0.5°C and four of them are 0.75°C or more, showing that duplicates alone can have the same or more impact as external factors.

Overall, other than the *Exact fragments* and *Similar fragments* categories, the majority of the remainder has differences of GC content and melting temperature of over 0.1°C. Many studies report these values to three digits of precision, or even more (57-62). The presence of duplicates means that these values in fact have considerable uncertainty. The impact depends on which duplicate type is considered. In this study, duplicates under the *Exact fragments*, *Similar fragments,* and *Low-identity* categories have comparatively higher differences than other categories. In contrast, *Exact sequences* and *Similar sequences* have only small differences. The impact of duplicates is also dependent on the specific organism: some have specific duplicate types with relatively large differences, and the overall difference is large as well; some only differ in specific duplicate types, and the overall difference is smaller; and so on. Thus it is valuable to be aware of the prevalence of different duplicate types in specific organisms.

In general, we find that duplicates bring much redundancy; this is certainly disadvantageous for studies such as sequence searching. Also, exemplars have distinct characteristics from their original groups such that sequence-based measurement involving duplicates may have biased results. The differences are more obvious for specific duplicate pairs within the groups. For studies that randomly select the records or have dataset with limited size, the results may be affected, due to possible considerable differences. Together they show that why de-duplication is necessary. Note that the purpose of our case study is not to argue that previous studies are wrong or try to better estimate melting temperatures. Our aim is only to show that the presence of duplicates, and of specific types of duplicates, can have a meaningful impact on biological studies based on sequence analysis. Furthermore, it provides evidence for the value of expert curation of sequence databases (63).

Our case study illustrates that different kinds of duplicates can have distinct impacts on biological studies. As described, the *Exact sequences* records have only a minor impact under the context of the case study. Such duplicates can be regarded as redundant. Redundancy increases the database size and slows down the database search, but may have no impact on biological studies.

In contrast, some duplicates can be defined as inconsistent. Their characteristics are substantially different to the ‘primary’ sequence record to which they correspond, so they can mislead sequence analysis. We need to be aware of the presence of such duplicates, and consider whether it they must be detected and managed.

In addition, we observe that the impact of these different duplicate types, and whether they should be considered to be redundant or inconsistent, is task-dependent. In the case of GC content analysis, duplicates under *Similar fragments* may have severe impact. For other tasks, there may be different effects; consider for example exploration of the correlation between non-codon and codon sequences (19) and the task of finding repeat sequence markers (20). We should measure the impact of duplicates in the context of such activities and then respond appropriately.

Duplicates can have impacts in other ways. Machine learning is a popular technique and effective technique for analysis of large sets of records. The presence of duplicates, however, may bias the performance of learning techniques because they can affect the inferred statistical distribution of data features. For example, it was found that much duplication existed in a popular dataset that has been widely used for evaluating machine learning methods used to detect anomalies (64); its training dataset has over 78% redundancy with 1,074,992 records over-represented into 4,898,431 records. Removal of the duplicates significantly changed reported performance, and behaviour, of methods developed on that data.

In bioinformatics, we also observe this problem. In earlier work we reproduced and evaluated a duplicate detection method (12) and found that it has poor generalisation performance because the training and testing dataset consists of only one duplicate type (52). Thus it is important to be aware of constructing the training and testing datasets based on representative instances. In general two strategies for addressing this issue: one using different candidate selection techniques (65); another is using large-scale validated benchmarks (66). In particular duplicate detection surveys point out the importance of the latter: as different individuals have different definitions or assumptions on what duplicates are, this often leads to the corresponding methods working only in narrow datasets (66).

## 6. Conclusion

Duplication, redundancy, and inconsistency have the potential to undermine the accuracy of analyses undertaken on bioinformatics databases, particularly if the analyses involve any form of summary or aggregation. We have undertaken a foundational analysis to understand the scale, kinds, and impacts of duplicates. For this work, we analysed a benchmark consisting of duplicates spotted by INSDC record submitters, one of the benchmarks we collected in (53). We have shown that the prevalence of duplicates in the broad nucleotide databases is potentially high. The study also illustrates the presence of diverse duplicate types and that different organisms have different prevalence of duplicates, making the situation even more complex. Our investigation suggests that different or even simplified definitions of duplicates, such as those in previous studies, may not be valuable in practice.

The quantitative measurement of these duplicate records showed that they can vary substantially from other records, and that different kinds of duplicates have distinct features that imply that they require different approaches for detection. As a preliminary case study, we considered the impact of these duplicates on measurements that depend on quantitative information in sequence databases (GC content and melting temperature analysis), which demonstrated that the presence of duplicates introduces error.

Our analysis illustrates that some duplicates only introduce redundancy, whereas other types lead to inconsistency. The impact of duplicates is also task-dependent; it is a fallacy to suppose that a database can be fully de-duplicated, as one task’s duplicate can be valuable information in another context.

The work we have presented based on the merge-based benchmark as a source of duplication, may not be fully representative of duplicates overall. Nevertheless, the collected data and the conclusions derived from them are reliable. Although records were merged due to different reasons, these reasons reflect the diversity and complexity of duplication. It is far from clear how the overall prevalence of duplication might be more comprehensively assessed. This would require a discovery method, which would inherently be biased by the assumptions of the method. We therefore present this work as a contribution to understanding what assumptions might be valid.

## Acknowledgements

We are grateful to Judice LY Koh and Alex Rudniy for explaining their duplicate detection methods. We also appreciate the database staff who have supported our work with domain expertise: Nicole Silvester and Clara Amid from EMBL ENA (advised on merged records in INSDC databases); Wayne Matten from NCBI (advised how to use BLAST to achieve good alignment results); and Elisabeth Gasteiger from UniProt (explained how UniProt staff removed redundant entries in UniProt TrEMBL).

Qingyu Chen’s work is supported by an International Research Scholarship from The University of Melbourne. The project receives funding from the Australian Research Council through a Discovery Project grant, DP150101550.

## Footnotes

1 http://www.uniprot.org/help/proteome_redundancy.

2 Quoted from Sylvain Poux, leader of manual curation and quality control in SwissProt.

3 http://www.uniprot.org/help/redundancy

4 http://www.uniprot.org/help/proteome_redundancy

5 http://insideuniprot.blogspot.com.au/2015/05/uniprot-knowledgebase-just-got-smaller.html

6 http://www.ncbi.nlm.nih.gov/nuccore/6017069?report=girevhist

7 http://www.ncbi.nlm.nih.gov/Taxonomy/taxonomyhome.html/

8 Ramona Britto and Benoit Bely, the key staff who removed over 45 million duplicate records from UniProtKB.

9 http://www.ncbi.nlm.nih.gov/books/NBK53704/

10 http://www.ebi.ac.uk/ena/submit/sequence-submission#how_to_update

11 http://www.ncbi.nlm.nih.gov/nuccore/56384585

